# Environmental conditions of recognition memory testing induce neurovascular changes in the hippocampus in a sex-specific manner in mice

**DOI:** 10.1101/2022.09.29.510187

**Authors:** Alice Cadoret, Laurence Dion-Albert, Sara Amrani, Laurianne Caron, Mathilde Théberge, Audrey Turmel, Manon Lebel, Caroline Menard

## Abstract

Experiences are linked to emotions impacting memory consolidation and associated brain neuronal circuits. Posttraumatic stress disorder is an example of strong negative emotions affecting memory processes by flashbacks of past traumas. Stress-related memory deficits are also observed in major depressive disorder (MDD). We recently highlighted that sex-specific blood-brain barrier (BBB) alterations underlie stress responses in mice and human depression. However, little is known about the relationship between emotional valence, memory encoding and BBB function. Here, we investigated the effects of novel object recognition (NOR) test, an experience considered of neutral emotional valence, on BBB properties in dorsal vs ventral hippocampus in the context of various environmental conditions (arena size, handling, age). The hippocampus is a brain area central for learning and memory processes with the dorsal and ventral subregions being associated with working memory vs reference memory retrieval, respectively. Expression of genes related to BBB integrity are altered in line with learning and memory processes in a region- and sex-specific manner. We observed correlations between poor learning, anxiety, stress-induced corticosterone release and changes in BBB-associated gene expression. Comparison of BBB transcriptomes between sexes also revealed profound differences at baseline in both ventral and dorsal hippocampus. Finally, we identified circulating vascular biomarkers, such as sE-selectin and Mmp-9, altered following NOR exposure supporting that recognition memory formation has an impact on the neurovasculature. Although deemed as a neutral valence test, NOR experimental conditions impact performance, highlighting the need to minimize anxiety when performing this commonly used test in mice.

**Significance Statement:** With this study, we aim to investigate the blood-brain barrier’s (BBB) role in memory acquisition and consolidation to unravel new mechanisms and decipher the involvement of non-neuronal cell types in these processes. For this purpose, male and female mice were subjected to a recognition memory test associated with a neutral emotional experience and impact on the transcriptomic profile of the BBB along with blood vascular biomarkers were evaluated under various experimental conditions. Crossing the BBB remains an important challenge to develop therapeutic drugs including in the context of memory deficits driven by psychiatric disorders or neurodegenerative diseases and thus, the possibility to directly target this barrier by better understanding its biology is attractive and innovating.

## Introduction

The BBB is a selectively permeable structure formed by pericytes, astrocytes, and endothelial cells sealed by tight junction proteins, which serves to prevent potentially harmful signals in the blood like blood cells, inflammatory cytokines, and pathogens, from entering the brain (Daneman and Prat, 2015; Keaney and Campbell, 2015; Menard et al., 2017b; Sweeney et al., 2019; Doney et al., 2022). Some cytokines, such as proinflammatory interleukin-6 (IL-6) and IL-1β, can cross the BBB via saturable transport (Banks et al., 1994; Banks et al., 1995; Dion-Albert et al., 2022a; Doney et al., 2022) to act directly on glial cells and neurons, affecting normal physiological processes such as temperature regulation, neuronal differentiation and survival, astrocyte proliferation and modulation of pain (Gadient and Otten, 1997).

Emotional processing and memory encoding can have a negative or positive valence, for example when associated with fear or reward. These processes involve various brain areas including the prefrontal cortex, hippocampus (HIP), nucleus accumbens and amygdala (Phillips et al., 2003), which, while closely interconnected, are each responsible for unique behaviors. For example, dense connections between the nucleus accumbens, a key hub in striatal reward circuitry, and HIP suggest that the HIP strengthens memory encoding based on the valence of a stimulus (Russo and Nestler, 2013). We reported that in male mice the BBB in the nucleus accumbens is more vulnerable to stress-induced immune response (Menard et al., 2017b) suggesting that negative emotional experience may have region-specific long-lasting effects on the neurovasculature. In fact, BBB breakdown is an early event in the aging human brain and cerebrovascular dysfunction begins in the HIP likely contributing to age-related cognitive decline (Montagne et al., 2015). While cerebrovascular dysfunction and immune response have been extensively studied in pathological conditions, little is known about their involvement in normal physiological processes including positive, neutral, and negative memory encoding.

Fear conditioning is a commonly used behavioral paradigm to explore the mechanisms of aversive learning and memory because it induces a well-defined response to a specific environmental stimulus (Johansen et al., 2011). Conversely, social interactions and food can be used as positive reinforcement in laboratory animals as they are essential for emotional well-being and act as incentive reward for maze learning (Trezza et al., 2011). The novel object recognition (NOR) test, which is based on the natural preference for novel objects displayed by rodents, is not associated with an emotional response (Antunes and Biala, 2012) and is thus considered of neutral valence. We took advantage of this last behavioral paradigm to compare the neurovascular biology underlying memory formation in different environmental contexts.

Sex differences in emotion and memory processes have been attributed to effects of female sex hormones (ter Horst et al., 2012). For example, female rodents are less anxious in a novel environment than males (Tropp and Markus, 2001). Males generally perform better in spatial tasks than females (ter Horst et al., 2012) but the female estrous cycle can bonify memory processing. Indeed, proestrus female rats are superior in learning the eye-blink conditioning task when compared to males, but not females in other estrous phases (Dalla et al., 2009; Dalla and Shors, 2009). Sex differences in the BBB may be implicated in emotional processing since sex hormones are potent modulators of neurovascular integrity (Dion-Albert et al., 2022a). We recently reported that vascular and BBB-related changes underlie stress susceptibility vs resilience in female mice (Dion-Albert et al., 2022b). Thus, in this study, both male and female mice were included to potentially unravel sex-specific BBB adaptations to memory experience.

Here, we combine behavioral studies performed in various experimental conditions (arena size, handling to reduce anxiety, age) with molecular profiling of BBB-related genes and measurement of blood circulating hormones and vascular biomarkers to characterize the impact of a neutral memory experience on the brain neurovasculature. Considering its crucial role in memory formation, investigation was centered on the HIP (Bannerman et al., 2014).

## Material and Methods

### Mice

Male and female C57BL/6 mice of 6-7 weeks or 6 months of age were purchased from Charles River and allowed 1 week of acclimatation. Mice were singled housed in a 12h/12h light/dark cycle. Food and water *ad libitum* were provided. All mouse procedures were performed in accordance with the Canadian Council on Animal Care (1993) as well as Université Laval animal care committees (#2022-1061).

### Novel object recognition (NOR)

One week after blood samples collection, experimental mice were divided into handling and non-handling groups, where that handling group was held 1 min per day for 7 consecutive days prior to the task. Naive control mice were also used for blood and molecular profiling, but they were never exposed to the arena or behavioral room. Dimensions of the arenas used were either 30×30×30cm or 50×50×50cm (as mentioned on Figures) and made with white Plexiglass. On the first day of the NOR task, mice were introduced to the empty arena for 5 minutes to acclimate them to it and to discriminate the novelty aspect of a novel environment (Ennaceur and Delacour, 1988; Leger et al., 2013; Lueptow, 2017). The second day, two novel identical objects were placed in opposite corners of the arena in equal distance from the walls. Mice were given 5 minutes to interact with the objects. One hour later, one of the objects, called the familiar, was kept in the arena at the same spot and the second one was replaced for a novel object. On the third day, 24 hours after the last test, the novel object was again replaced by a novel one and it was repeated 5 min later. Mice were sacrificed 2 hours after the last test. Experiments were conducted under red light and animals were moved in the behavioral room 1 hour before the first test each day to acclimatize them. Arena was cleaned with a hydrogen peroxyde solution (Prevail) at the beginning, between each mouse and at the end of the trials. Objects were different enough to be discriminated by mice but had similar degree of complexity (texture, shape, etc.) to minimize any object preference that could bias the results.

### Behavioral analysis

Video sessions were collected from behavioral experiments and analyzed with the video-tracking analysis program Ethovision XT (Noldus Information Technology). Nose-point detection was used to calculate the time mice interacted with the objects in a determined zone of 2,5 centimeters around the objects. NOR ratio was calculated by dividing time spent with the novel object on the total time spent with both objects, for the 5 minutes, 1 hour and 24 hours timepoints.

### Tissue collection and gene expression

Dorsal and ventral hippocampus (HippD and HippV) samples were collected and processed as described previously (Golden et al., 2013; Menard et al., 2017b). Bilateral 2.0mm punches were collected from 1-mm coronal slices on wet ice after rapid decapitation and immediately placed on dry ice and stored at −80 °C until use. RNA was isolated using TRIzol (Invitrogen) homogenization and chloroform layer separation. The clear RNA layer was processed using the Pure Link RNA mini kit (Life Technologies) and analyzed with NanoDrop (Thermo Fisher Scientific). RNA was reverse transcribed to cDNA with Maxima-H-minus cDNA synthesis kit (Fisher Scientific) and diluted to 500 μL. For each qPCR reaction, 3 μL of sample cDNA, 5 μL of Power up SYBR green (Fisher Scientific), 1 μL PrimeTime® qPCR primer (Integrated DNA Technologies) and 1 μL ddH20 was added to each well. Samples were heated to 95°C for 2 min, followed by 40 cycles of 95°C for 15 s, 60°C for 33 s and 72°C for 33 s. Analysis was done using the ΔΔCt method and samples were normalized to the Gapdh mouse housekeeping gene. Primer pairs (Integrated DNA Technologies) are listed in Supplementary Table 1.

### Corticosterone (CORT) measurement from serum samples

Blood samples were collected 1 week before the start of NOR protocol and during brain tissue collection. Blood was allowed to clot for at least 1 h before being centrifuged at 10,000 RPM at RT for 2 min. The supernatant was collected and spun again at 3000 RPM for 10 min. The clear supernatant (serum) was collected, aliquoted and stored at −80°C until use. Serum CORT levels were determined using the DetectX CORT enzyme immunoassay (Arbor Assays, Ann Arbor, MI). (Protocol listed on the product details link). Data was reduced against a four-parameter logistic curve using the Gen 5.0 software and samples with a coefficient of variation above 15% were removed from the analysis.

### Milliplex Assays for serum samples

MILLIPLEX Map 96-Well Plate Assay from Millipore Sigma (mouse CVD magnetic panel MCV1DMAG-77K-07 and Mouse Angiogenesis/GF MAG – MAGPMAG-24K-18) were assessed following protocol guidelines (EDM Millipore, MA, USA). Briefly, plate was prepared with 200 uL Assay Buffer on a plate shaker for 10min RT. 25 uL of standards and controls were added on the plate, then 25 uL of Samples (diluted 1:2 for the MAGPMAG-24K plate and 1:20 for the MCVD1MAG-77K) (Dion-Albert et al., 2022b). Beads mix was added, and after the plate was sealed and put on a plate shaker at 600 rpm overnight at 4°C. Next, the plate was washed multiple times, followed by addition of 25 uL of Detection Antibody for 1 hour at RT and then, 25 uL of Streptavidin-Phycoerythrin for 30 minutes (RT). The plate was washed several times, resuspended with Sheath Fluid for 5 minutes on a plate shaker, then read with the Bio-Plex 200© plate reader (Bio-Rad Laboratories, ON, Canada). Data was reduced against a five-parameter logistic curve using the Bio-Plex Manager software and samples with a coefficient of variation above 15% or Out of Range of the Standard curve were removed from the analysis.

### Statistic Analysis

All data were processed for statistical analysis with Graph Pad Prism software. Outliers were identified by Grubbs’ test with significance level of Alpha= 0.05. Behavioral data were analysed by using two-way ANOVA and assessed for multiple comparisons with Bonferroni post-hoc analysis. Gene expression profiles were analysed by using unpaired t-test between naive mice vs NOR mice fold changes or through two-way ANOVA followed by multiple comparisons with Bonferroni to compare environmental conditions or sexes. Relationship between BBB-related gene expression and NOR ratio or CORT level was analyzed with Pearson correlations. Corticosterone serum levels were analysed by using two-way ANOVA followed by multiple comparisons with Bonferroni post-tests. Statistical significance was set at *p* < 0.05 with \**p* < 0.05; \*\**p* < 0.01; \*\*\**p* < 0.001; \*\*\*\**p* < 0.0001. Values between *p* = 0.05 and *p* = 0.10 were considered as trending without reaching significance.

## Results

### Object recognition memory testing in a large arena alters behavioral performance along with neurovascular gene expression in the hippocampus

The NOR test is commonly used to investigate learning and memory in mice and is based on rodent’s natural behavior to explore novelty in their environment (Lueptow, 2017). First, 8-week-old C57Bl/6 male mice were subjected to the NOR paradigm consisting of a session of habituation to the empty arena on day 1, a training session with two similar objects on day 2 followed by three trials with a novel object at 5 min, 1h, or 24h before tissue collection (**Fig. 1A**). Blood was drawn prior and after the NOR paradigm to evaluate changes induced by the test in circulating stress-related hormone CORT. With a 50×50 cm arena, we observed NOR ratio, as calculated by the time spent with the novel vs familiar object, above 0.5 for 70% and 66% of the mice at 5 min and 24h, respectively, suggesting overall preference for the novel object (**Fig.1B**, left). However, the time spent with the objects was generally low (less than 10 sec, **Fig.1B**, right) suggesting high anxiety. A 2^nd^ cohort of mice was subjected to the same NOR paradigm but in a smaller 30×30 cm arena. It did not alter NOR ratio (**Fig.1B**, left) however, time spent with the objects was significantly increased (**Fig.1B**, right, two-way ANOVA arena effect: \*\*\**p=0*.*0003*). Stress exposure can alter BBB integrity (Menard et al., 2017b) including in the HIP (Santha et al., 2015) which is central for memory encoding. Thus, we investigated expression of several BBB-related genes including growth factors (*Bdnf:* Brain-derived neurotrophic factor, *Fgf2:* Fibroblast growth factor 2, *Vegfa:* Vascular endothelial growth factor), tight junction proteins (*Cldn5*: Claudin-5, *Ocln*: Occludin, *Tjp1*: Tight junction protein 1), astrocyte glial fibrillary acidic protein (*Gfap*) and platelet endothelial cell adhesion molecule (*Pecam1*) in the ventral and dorsal HIP. The dorsal HIP is responsible for cognitive processing such as spatial learning, while the ventral HIP is associated with emotional processing such as anxiety (Moser et al., 1993; Bannerman et al., 2003; Bannerman et al., 2014; Hauser et al., 2020). Decreased expression of *Cldn5* (\**p=0*.*0235*), *Ocln* (\**p=0*.*0462*), *Tjp1* (\**p=0*.*0160*), *Gfap* (\**p=0*.*0354*) and *Pecam1* (\**p=0*.*0234*) was observed in the ventral HIP of male mice that performed the NOR test in the 50×50 cm arena when compared to naive animals **(Fig.1C)**. In contrast, an increase in *Fgf2* (\*\**p=0*.*0033*) was measured for the mice in the 30×30 cm arena **(Fig.1C)** indicating that the size or the arena has a direct impact not only on object recognition memory performance but also on transcription in the ventral HIP neurovasculature. Direct statistical comparison confirmed a significant effect of the arena for *Fgf2* (**Fig.1D**, top left, two-way ANOVA arena effect: \*\**p=0*.*0031*) and *Gfap* (**Fig.1D**, bottom right, two-way ANOVA arena effect: \**p=0*.*0325*) driven by the mice exposed to the NOR paradigm (Sidak’s multiple comparisons test: \*\*\**p=0*.*0005* for *Fgf2* and \**p=0*.*0224* for *Gfap*). As for the dorsal HIP, changes in gene expression were noted only for the 50×50 cm arena with an increase in *Bdnf* (\*\**p=0*.*0069*), *Fgf2* (\**p=0*.*0465*), *Vegfa* (\*\**p=0*.*0025*), *Ocln* (\*\**p=0*.*0057*) and *Tjp1* (\*\**p=0*.*0019*) for male mice exposed to the NOR paradigm vs naive animals **(Fig.1E)**. Direct statistical comparison revealed a significant effect of the arena for *Fgf2* again in this HIP subregion (**Fig.1F**, top left, two-way ANOVA arena effect: \**p=0*.*0222*) as well as *Vegfa* (**Fig.1F**, top right, two-way ANOVA arena effect: \**p=0*.*0161*). Like for the ventral HIP, alterations were driven by the mouse groups subjected to the NOR test (Sidak’s multiple comparisons test: \**p=0*.*0120* for *Fgf2* and \*\**p=0*.*0054* for *Vegfa*). To confirm that the larger arena is inducing anxiety, CORT level was compared prior vs after exposure to the NOR paradigm. As expected, a significant increase in circulating CORT was observed after the test (**Fig.1G**, two-way ANOVA before vs after effect: \*\**p=0*.*0031*), however, this increment was much lower for the 30×30 cm arena group (Tukey’s multiple comparison test: \*\*\**p=0*.*0007*). Blood CORT level is correlated negatively with the expression of tight junction *Tjp1* in the dorsal, but not ventral, HIP (**Fig.1H**). Overall, our results suggest that performing the NOR test in a larger area promotes anxiety altering neurovascular gene expression in the HIP in a subregion-specific manner.

**Figure 1.**
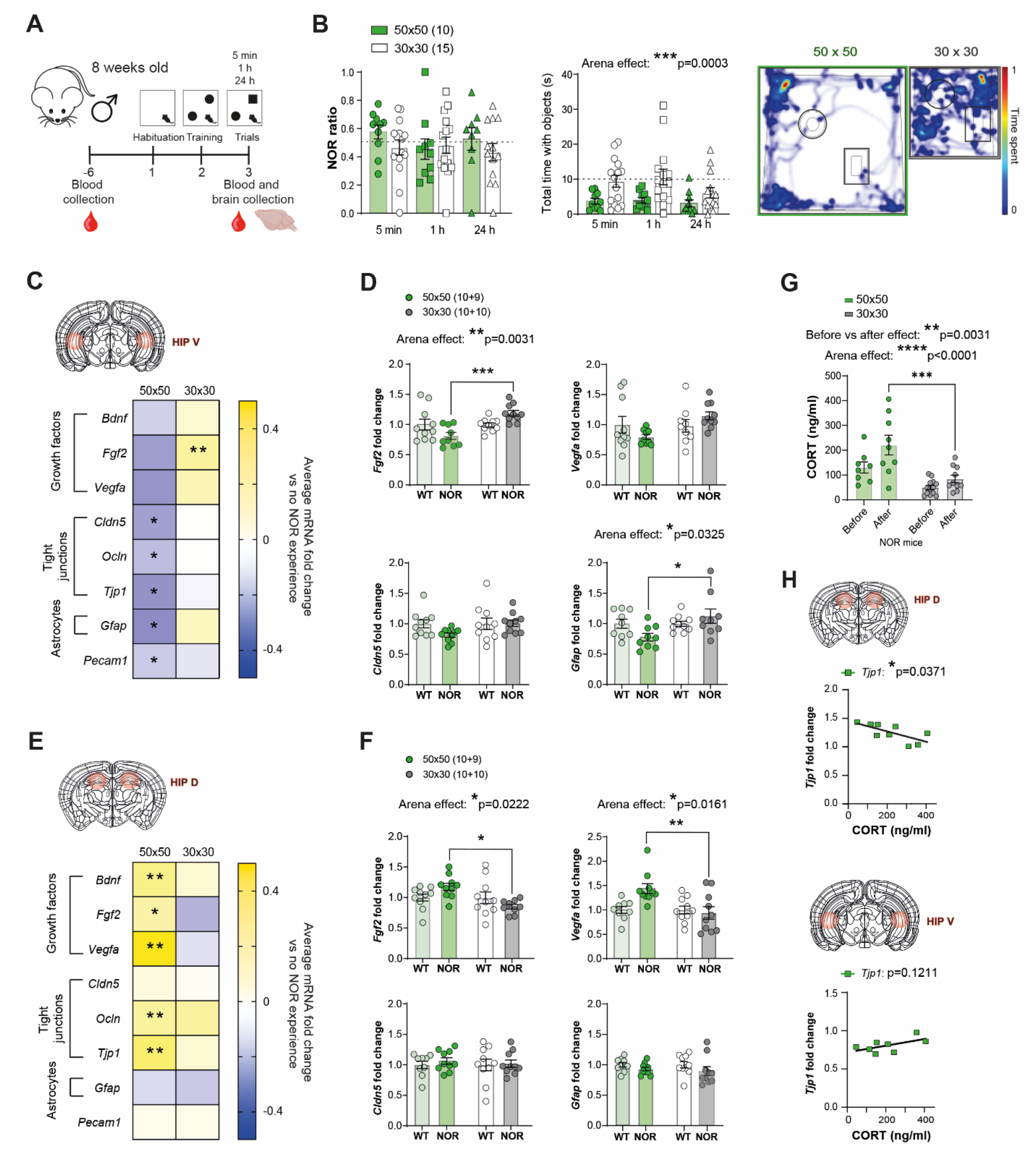
Object recognition memory testing in a large arena alters performance and neurovascular expression in the ventral and dorsal hippocampus. **(a)** Experimental behavioral NOR performances at 5 minutes, 1 hour and 24 hours time points are influenced by arena size when looking at **(b)** the time interacting with objects but not when looking at the NOR ratio (2-way ANOVA). **(c)** Quantitative PCR in HippV reveals tight junction genes expression decreased in mice that went through NOR test in 50×50 arena vs naive mice, but no changes are observed for 30×30 NOR mice (t-test). **(d)** *Fgf2, Vegfa* and *Gfap* expression go in opposite direction for expression in NOR mice 50×50 vs 30×30, as *Cldn5* expression decreased in 50×50 but not in 30×30 NOR mice (t-test and 2-way ANOVA). **(e)** In HippD, some vascular and BBB gene are up regulated in the 50×50 NOR mice cohort but not in the 30×30 mice, **(f)** with *Fgf2* and *Vegfa* going in opposite direction in mRNA expression, and no change at all for *Cldn5* and *Gfap*. **(g)** Corticosterone (CORT) serum levels are higher before and after in NOR mice that realized the test in 50×50 arena (Arena effect, 2-way ANOVA). CORT levels seem to be correlated with HippD *Tjp1* expression in the 50×50 NOR mice cohort (Pearson correlation). Data represent mean ± SEM with the number of animals indicated on legends and graphs by individual data points; *p ≤ 0.05, **p ≤ 0.01, ***, p ≤ 0.001, ****p ≤ 0.0001.

### Handling improves object recognition memory and induces changes in neurovascular gene expression mostly in the ventral hippocampus

Handling is known to reduce stress in rodents (Marcotte et al., 2021). We evaluated if handling prior to the NOR paradigm could further the improvement observed after reduction of the arena size (**Fig.1**). 8-week-old C57Bl/6 male mice were handled every day for 7 days prior NOR testing (**Fig.2A**) as described in the Methods. No significant change was observed between groups for the NOR ratio (**Fig.2B**, left), however, the handled mice spent more time interacting with the objects (**Fig.2B**, right, two-way ANOVA handling effect: \*\*\*\**p<0*.*0001*) particularly for the 5-min time point (Bonferroni’s multiple comparisons test: \*\**p=0*.*0018*). Next, changes in neurovascular gene expression were explored (**Fig.2C**) revealing an increase in *Fgf2* (\*\**p=0*.*0033*) in the ventral HIP when non-handled mice were compared to naive animals. Mouse NOR performances were uneven at the various time points tested (5 min, 1h, 24h) so comparisons were done with gene expression for each (**Fig.2D**). In contrast to arena size, handling had no significant effect on BBB-related genes of the dorsal HIP (**Fig.2E-F**). Nevertheless, this manipulation reduced circulating CORT (**Fig.2G**, two-way ANOVA before vs after effect: \*\**p=0*.*0023*) even in the group of mice not exposed to the NOR paradigm (handling effect: \*\**p=0*.*0075*) confirming anxiolytic effect. Blood CORT was again negatively correlated with the expression of tight junctions, this time *Cldn5*, in the ventral HIP (*p=0*.*0651*) (**Fig.2H**). Moreover, a change was observed for both hippocampal subregions for the astrocytic marker *Gfap* (\**p=0*.*0256* for the ventral HIP and *p=0*.*0601* for the dorsal HIP). Altogether, these results reinforce the need to consider handling prior to NOR memory testing to prevent stress-induced BBB alterations.

**Figure 2.**
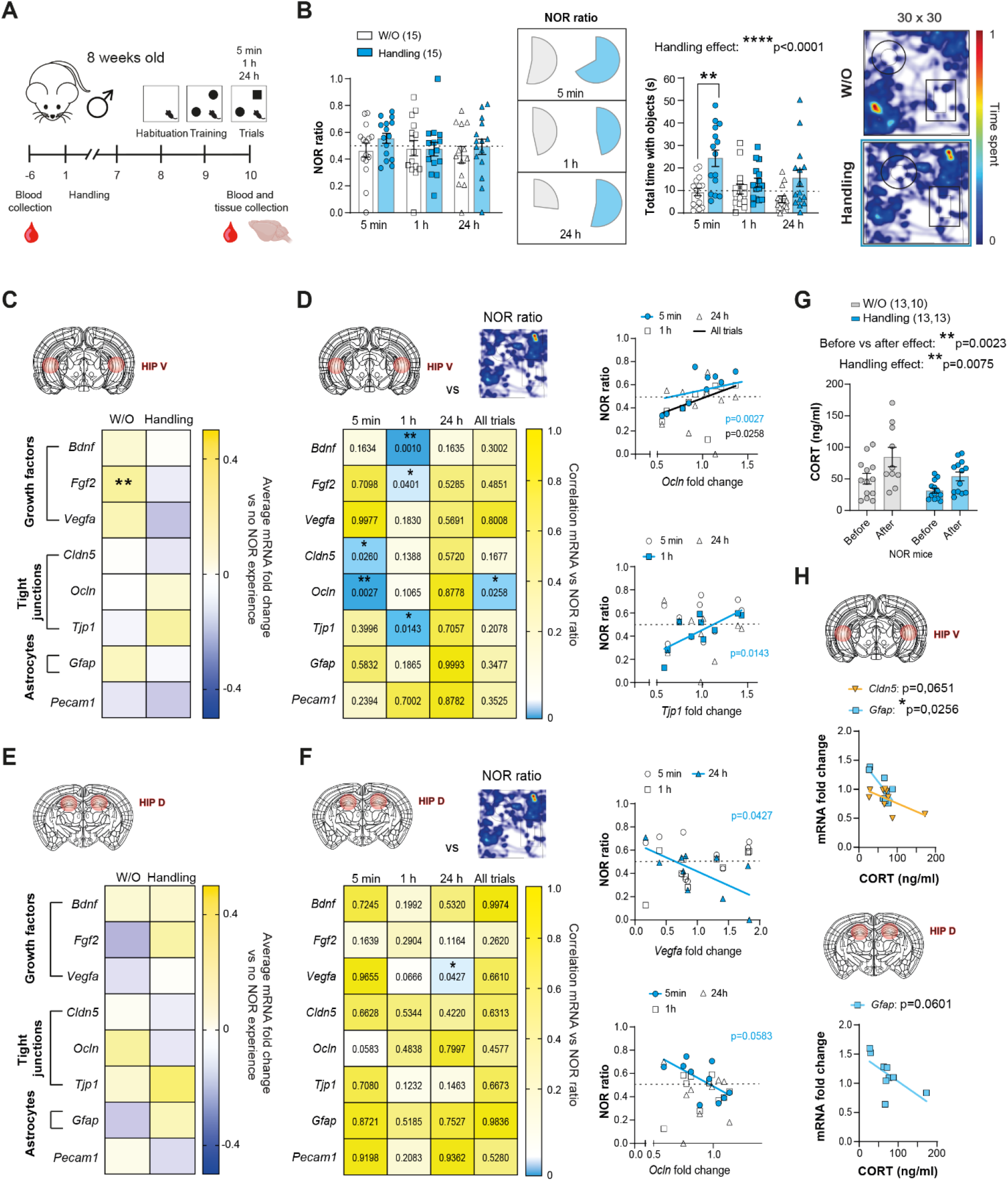
Handling improves object recognition memory and induces changes in neurovascular gene expression mostly in the ventral hippocampus. **(a)** Experimental behavioral NOR testing with added handling for one cohort prior to the NOR testing. **(b)** Proportion of NOR ratio and time interacting with objects is increased in NOR mice that were handled compared to NOR mice that were not (2-way ANOVA). **(c)** Quantitative PCR reveals few changes in BBB and vascular gene expression in HippV, **(d)** but few genes such as *Ocln* and *Tjp1* expressions are correlated with memory performance indicator in different NOR test time points (Pearson correlation). **(e)** Gene expression is not impacted following a NOR test in HippD of mice vs naive controls and **(f)** slightly correlated with NOR ratio. **(g)** Corticosterone (CORT) serum levels are lower before and after in handled NOR mice (Handling effect, 2-way ANOVA). **(h)** *Gfap* expression in HippV and HippD seems to tend to be correlated with CORT serum levels, as *Cldn5* in HippV (Pearson correlation). Data represent mean ± SEM with the number of animals indicated on legends and graphs by individual data points; *p ≤ 0.05, **p ≤ 0.01, ***, p ≤ 0.001, ****p ≤ 0.0001.

### Age had minimal effect on novel object recognition memory performance and neurovascular gene expression despite a strong increase in circulating CORT

Stress-related paradigms revealing BBB dysfunction were performed in young adults (8-12 weeks) (Menard et al., 2017b; Dion-Albert et al., 2022b) while learning and memory studies are often conducted on older adult rodents (6+ months)(Menard et al., 2013; Menard et al., 2014). Thus, we next evaluated whether age-specific neurovascular changes could underlie recognition memory performance during adulthood. 6-month-old C57Bl/6 male mice were handled every day for 7 days prior to NOR testing (**Fig.3A**). Age had no impact on memory performance (**Fig.3B**, two-way ANOVA age effect: *p=0*.*6449* for NOR ratio and *p=0*.*1033* for time with objects). Accordingly, no differences in growth factors or tight junction gene expression were noted between 8-week-old and 6-month-old that experienced the NOR paradigm **Fig.3C**); however, the astrocyte marker *Gfap* (\*\*\*\**p<0*.*0001*) and endothelial marker *Pecam1* (\*\*\**p=0*.*0003*) were decreased in the ventral HIP of older mice. When compared to their naive counterparts, male mice exposed to the NOR test had lower *Cldn5, Gfap* and *Pecam1* and this effect was driven by older adults (**Fig.3D**). Conversely, no change was observed for the dorsal HIP (**Fig.3E-F**), suggesting that the neurovasculature of HIP subregions are differentially impacted by aging. As in previous cohorts, blood circulating CORT level was compared prior vs after the NOR paradigm. A strong increase was measured in older adults (**Fig.3G**, two-way ANOVA before vs after effect: \*\*\*\**p<0*.*0001*) but also when only age was considered as a variable (**Fig.3G**, \*\*\*\**p<0*.*0001*). This could be related to the loss of *Cldn5, Gfap* and *Pecam1* in the ventral HIP (**Fig.3C-D**) since compensatory changes appear to be present in the dorsal HIP with a trend observed for an increase in the tight junction *Ocln* despite elevated circulating CORT **Fig.3H**, *p=0*.*0524*). Our results suggest that NOR memory performance is maintained throughout adulthood even as BBB-related alterations emerge in the ventral HIP of older animals and sensitivity of the HPA axis increases.

**Figure 3.**
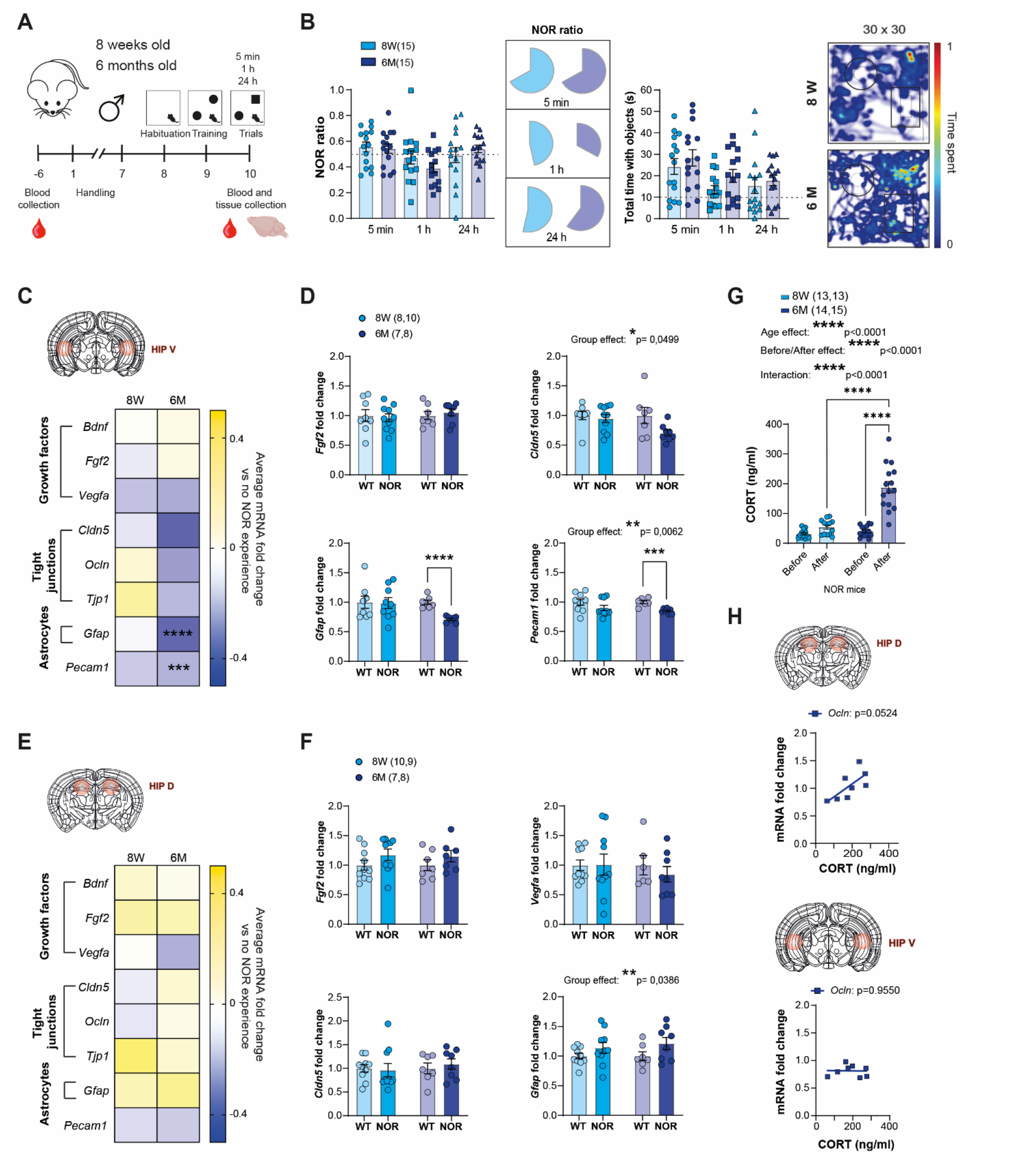
Age has minimal effect on novel object recognition performance and neurovascular gene expression despite a strong increase in circulating CORT. **(a)** Experimental behavioral NOR testing for 8 weeks and 6 months old cohorts for the NOR testing. **(b)** Proportion and mean of NOR ratios and total exploratory time with objects are not changed either for 8 weeks or 6 months old mice (2-way ANOVA). **(c)** Quantitative PCR reveals **(d)** decreased *Gfap* and *Pecam1* expression in HippV for 6 months old mice after NOR test, but not in the HippV of 8 weeks old, when compared with naive mice of corresponding age (unpaired t-test). **(e)** In HippD, no BBB and vascular gene expressions are different between both ages, and interestingly **(f)** *Gfap* expression seems to be increased with a NOR effect in NOR mice for both ages, but no other genes. **(g)** Corticosterone (CORT) serum levels are increased in a tremendous way for 6 months old NOR mice after the NOR test compared to before and to 8 weeks old after the test (2-way ANOVA), with a strong effect of age and time point. **(h)** 6 months old NOR mice show CORT levels that correlate with *Ocln* expression fold change in HippD (Pearson correlation). Data represent mean ± SEM with the number of animals indicated on legends and graphs by individual data points; *p ≤ 0.05, **p ≤ 0.01, ***, p ≤ 0.001, ****p ≤ 0.0001.

### Males and females perform the NOR test similarly but show differences in neurovascular gene expression and circulating CORT level

Stress as well as sex hormones can have an impact on memory processes (ter Horst et al., 2012). Thus, we next subjected 8-week-old female C56Bl/6 mice to handling followed by the NOR test paradigm (**Fig.4A**). Both NOR ratio (two-way ANOVA sex effect: *p=0*.*4177*) and time spent with objects (*p=0*.*8697)* were comparable between sexes (**Fig.4B**). Males and females are characterized by major biological differences such as sex chromosomes and different level of gonadal hormones but can behave alike. This observation led to the theory that to reach a convergent behavioral endpoint and overcome sex differences, the brain exploited compensatory physiological mechanisms (De Vries, 2004; McCarthy et al., 2012; Bangasser and Wicks, 2017). We thus compared the impact of NOR testing on BBB-related genes in the ventral and dorsal HIP. No difference was noted between males and females exposed to NOR in the ventral HIP (**Fig.4C**), however, in female mice *Vegfa* (\*\**p=0*.*005*), *Ocln* (\*\**p=0*.*0016*) and *Tjp1* (\*\*\**p=0*.*0009*) correlated significantly with NOR ratio (**Fig.4D**). In the dorsal HIP, *Bdnf* expression increased in females only (**Fig.4E**, unpaired t-test: \*\**p=0*.*0096*) with no correlation between NOR ratio and tight junction expression for this subregion (**Fig.4F**). Analysis of blood CORT revealed elevated baseline level of this hormone in females when compared to males (**Fig.4G**, two-way ANOVA sex effect: \*\*\*\**p<0*.*0001*). Sex differences in circulating CORT remain after NOR testing (**Fig.4G**, two-way ANOVA interaction sex x NOR: \*\*\*\**p<0*.*0001*). In contrast to males, CORT does not seem to directly impact BBB-associated genes in females (**Fig.4H**), highlighting the importance of considering sex as a biological variable while investigating the biological mechanisms underlying learning and memory processes.

**Figure 4.**
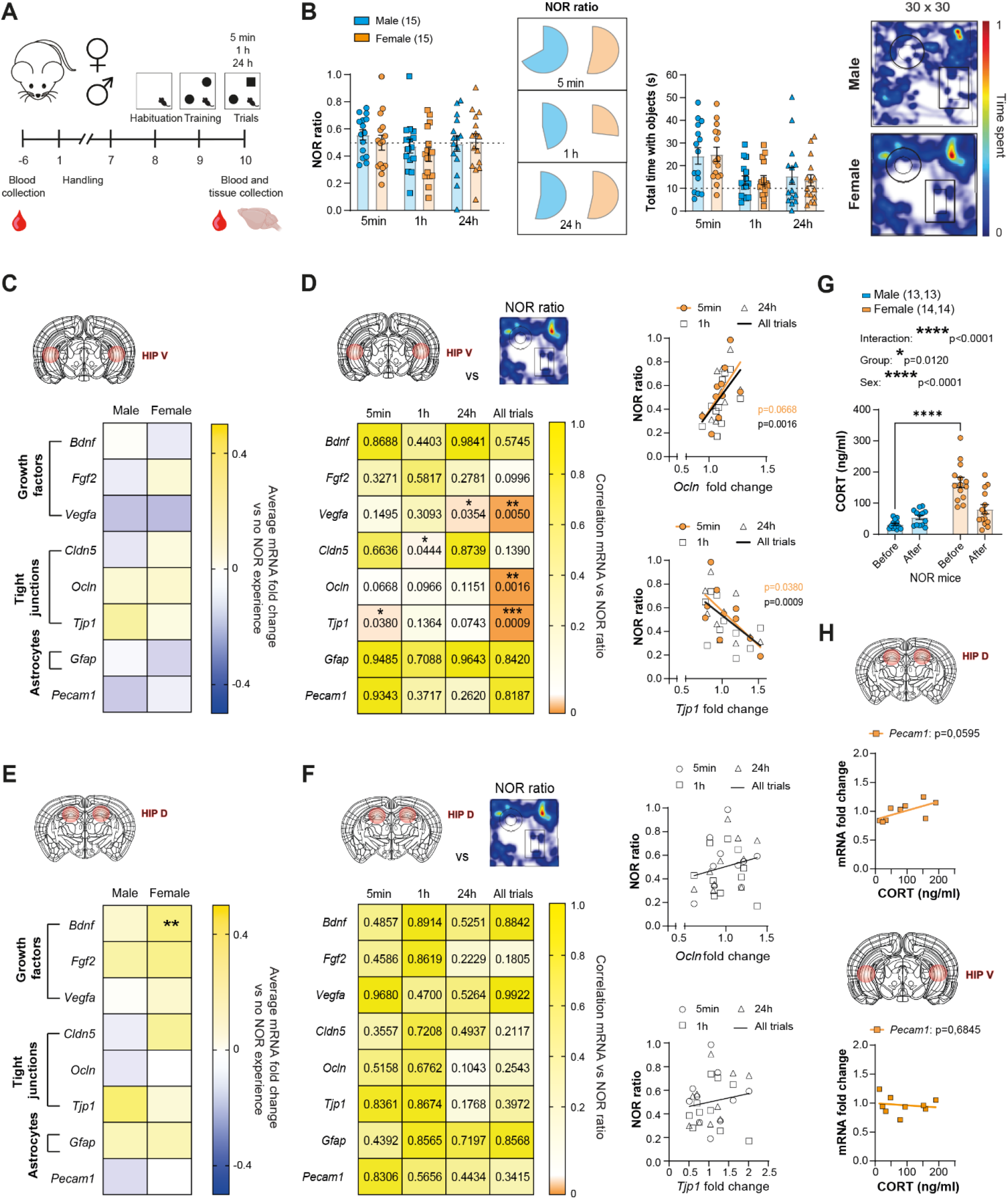
Males and females performed similarly nevertheless, sex differences were observed for neurovascular gene expression and level of circulating CORT. **(a)** Experimental behavioral NOR testing for male and female cohorts for the NOR testing. **(b)** Proportion and mean of NOR ratios and total exploratory time with objects are not changed either for male or female mice (2-way ANOVA). **(c)** Quantitative PCR reveals no differences in HippV gene expression between male and female NOR mice, **(d)** but presents *Vegfa, Cldn5, Ocln, Tjp1* expression levels that correlate with NOR performance for different time points. **(e)** In HippD, no BBB and vascular gene expressions are different between sexes, except *Bdnf* which is upregulated in female NOR mice (unpaired t-test). **(f)** No correlation between vascular and BBB gene expression and NOR ratio are observed in HippD of female NOR mice (Pearson correlation). **(g)** Corticosterone (CORT) serum levels are higher in females NOR mice before the test, to come back to a similar level than male NOR mice after the test (2-way ANOVA). **(h)** In HippD, *Pecam1* expression is going toward a correlation with CORT serum levels (Pearson correlation). Data represent mean ± SEM with the number of animals indicated on legends and graphs by individual data points; *p ≤ 0.05, **p ≤ 0.01, ***, p ≤ 0.001, ****p ≤ 0.0001.

### Baseline sex and regional differences in neurovascular gene expression

Our recent work exposed sex-specific differences in transcriptomic profile for BBB-related genes in the prefrontal cortex and nucleus accumbens of male and female mice even at baseline (Dion-Albert et al., 2022b). Sex differences in the brain vasculature are understudied hence, we explored if they could be present in the ventral and dorsal HIP as well. Strikingly, expression of BBB tight junctions *Cldn5* (\*\*\**p=0*.*0004*), *Ocln* (\*\*\*\**p<0*.*0001*), *Tjp1* (\*\*\*\**p<0*.*0001*), growth factors *Bdnf* (\*\*\*\**p<0*.*0001*), *Fgf2* (\**p=0*.*0368*), *Vegfa* (\*\*\**p=0*.*0002*), astrocytic *Gfap* (\*\*\**p=0*.*0005*) and endothelium *Pecam1* (\*\**p=0*.*0021*) were all lower in the ventral HIP of naive female C57Bl/6 when compared to males (**Fig.5A**, left and **B**, top). Even when gene expression was normalized on *Pecam1* to rule out an overall difference in the neurovascular network, *Ocln* (\*\**p=0*.*0026*) and *Tjp1* (\*\*\*\**p<0*.*0001*) remained lowly expressed in the female ventral HIP vs their male counterparts (**Fig.5A**, right and **B**, bottom). As for the dorsal HIP, while BBB tight junctions *Cldn5* (\*\*\*\**p<0*.*0001*), *Ocln* (\**p=0*.*0214*), astrocytic *Gfap* (\**p=0*.*0194*) and endothelium *Pecam1* (\*\*\*\**p<0*.*0001*) expression were again lower in females, *Tjp1* was increased (\*\**p=0*.*0019*) and no sex difference was noted for growth factors (**Fig.5C**, left and **D**, top). *Pecam1* normalization did not change the findings for *Cldn5* (\*\*\**p=0*.*0008*) and *Tjp1* (\*\*\*\**p<0*.*0001*) but reversed the sex-specific profile for *Gfap* (\**p=0*.*0119*) (**Fig.5C**, right and **D**, bottom). Finally, baseline gene expression was directly compared between the HIP dorsal and ventral subregions. Most BBB-related genes and growth factors, except *Vegfa* for males, appears to be more expressed in the dorsal when compared to the ventral HIP (**Fig.5E-F**). Altogether, these results highlight the importance to consider sex as a biological variable when investigating hippocampus-related behaviors and underlying mechanisms.

**Figure 5.**
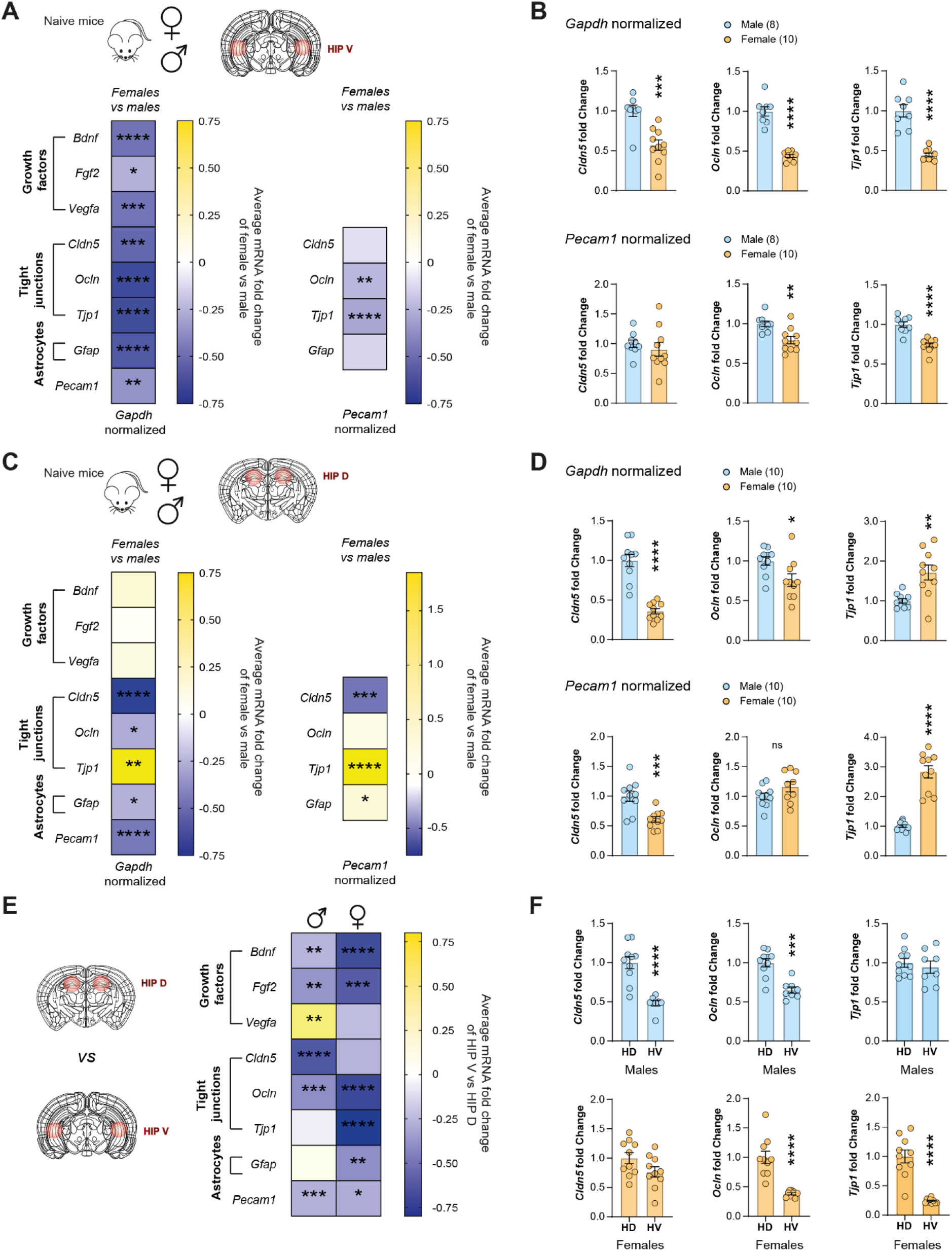
Baseline sex and regional differences of neurovascular gene expression. **(a)** When looking at naive mice that did not experienced any memory experience, we observed **(b)** downregulation of BBB and vascular gene expression from female mice brain when compared to male ones in the ventral hippocampus. (t-test) **(c)** This can be noticed as well for TJs genes (*Cldn5, Ocln* and *Tjp1*) when normalized on endothelial gene *Pecam1*. **(d)** In dorsal hippocampus, downregulation of basal gene expression is mostly observed in TJs, but not in growth factors when looking at female vs male mouse brain. **(e)** *Cldn5* is decreased in female brains, either normalized on *Gapdh* or *Pecam1*, on the other hand *Tjp1* is increased either normalized on *Gapdh* or *Pecam1* (t-test). **(f)** In male and female, when we compared gene expression levels of HippV compared on HippD, most of the vascular and BBB genes are downregulated on a basal level in the ventral region of the hippocampus, expect for *Vegfa* in male which is higher in HippV than in HippD (t-test). **(g)** In male mice, *Cldn5* and *Ocln* basal levels are decreased in HippV compared to HippD, as for *Ocln* in female mice, but not *Cldn5* (t-test). Data represent mean ± SEM with the number of animals indicated on legends and graphs by individual data points; *p ≤ 0.05, **p ≤ 0.01, ***, p ≤ 0.001, ****p ≤ 0.0001.

### NOR experimental conditions are associated with changes in circulating vascular markers

We recently reported that stress-induced BBB dysfunction is associated with sex-specific increase in blood vascular biomarkers (Dion-Albert et al., 2022b) providing an indirect measure of BBB health status. Thus, we investigated here if recognition memory performances in the various environmental conditions are also reflected in the circulation by taking advantage of commercially available Milliplex MAP mouse cardiovascular disease and mouse angiogenesis/growth factor magnetic bead panels. Blood was collected after exposure to the NOR test (**Fig.6A**) and then processed to measure numerous analytes: soluble adhesion molecule sE-Selectin, endothelium Pecam-1, soluble platelet sP-Selectin, matrix metallopeptidase 9 (Mmp-9), inflammation-related granulocyte-colony stimulating factor (G-Csf), soluble anaplastic lymphoma kinase (sAlk-1), fibroblast growth factor 2 (Fgf-2), hepatocyte growth factor (Hgf), interleukin (Il)-17A, chemokine (C-X-C motif) ligand 1 (CXCL1, or KC) and vascular endothelial growth factor (Vegf)-c. Intriguingly, running the NOR test in the small 30×30 arena (**Fig.1**) significantly reduced blood sE-selectin (\*\*\*\**p<0*.*0001*), sP-selectin (\**p=0*.*0202*) and Mmp-9 (\*\*\**p=0*.*0002*) while a trend was observed for G-Csf (*p=0*.*0618*) and Hgf (*p=0*.*0717*) (**Fig.6B**). Meanwhile, handling tends to reduce levels of Mmp-9 (*p=0*.*0796*), G-Csf (*p=0*.*0618*) and sAlk-1 (*p=0*.*0907*). No difference was noted for females when comparing NOR and naive cohorts (**Fig.6B**). Direct comparison of sE-selectin level between NOR cohorts performed in the various conditions showed a reduction for males who performed in the small arena (\*\**p=0*.*0072*) (**Fig.6C**, top left). Handling moreover increased sE-selectin blood level in males (\**p=0*.*0148*, **Fig.6C**, top middle) however, this effect was sex-specific (\**p=0*.*0177*, **Fig.6C**, top right) (*p=0*.*0907*). Similar comparisons were done for Mmp9, and the only significant difference was noted for the small vs larger arena (\**p=0*.*0249*). These results suggest circulating markers may reflect changes in the neurovasculature induced by learning and memory in various experimental conditions.

**Figure 6.**
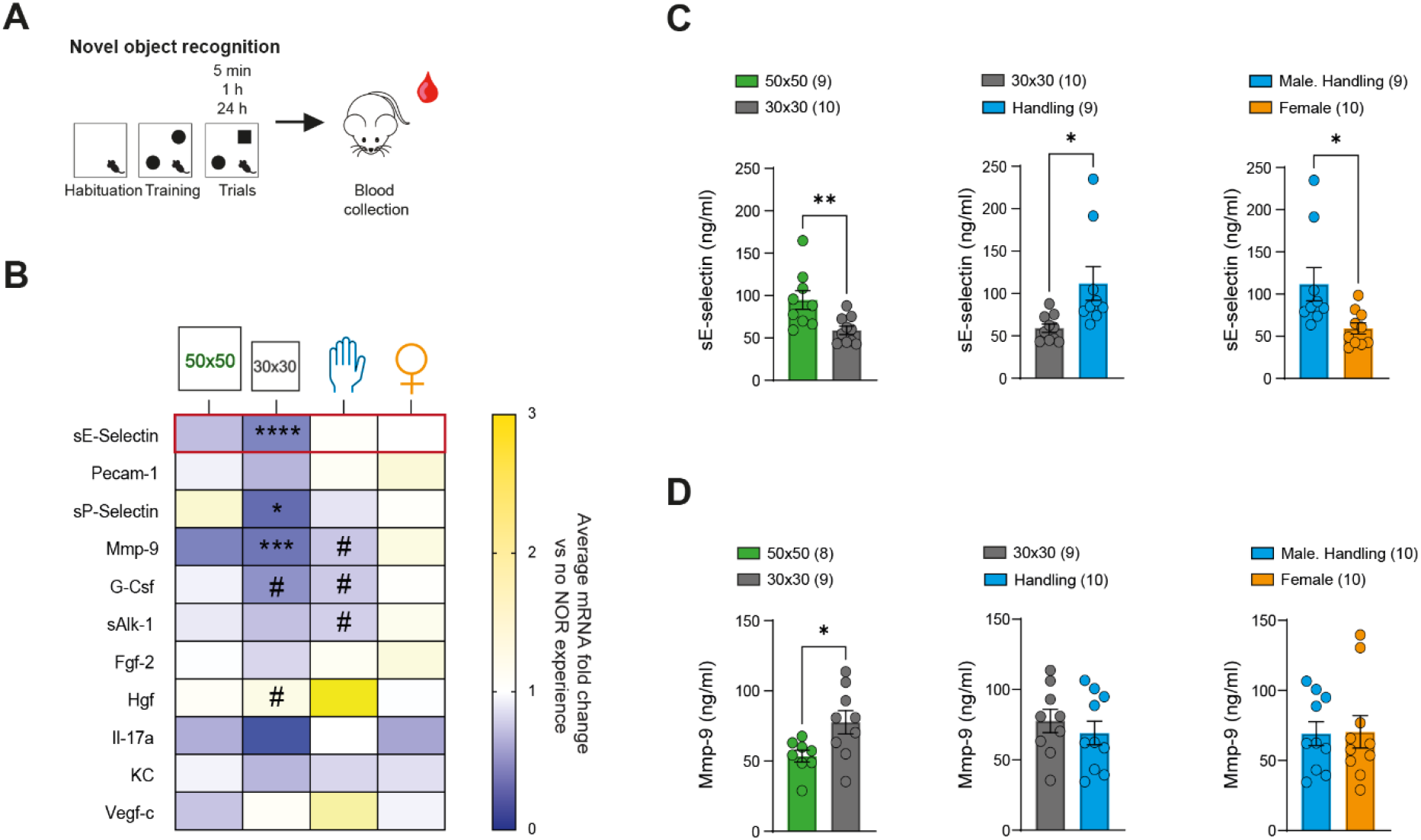
NOR conditions are associated with changes in circulating vascular biomarkers. **(a)** Blood is collected after NOR test 2-3h after the last trial. **(b)** In NOR cohorts’ vs naive mice, we observed lower circulating vascular markers, particularly for sE-selectin, sP-selectin, Mmp-9, G-Csf in the male 30×30 non-handled cohort and male handled cohort (t-test). **(c)** When all NOR cohorts were compared to each other, lower sE-selectin level was noted for the smaller 30×30 cm vs the large 50×50 arena. Handling increased circulating sE-selectin in males but not females. (**d**) In contrast to sE-selectin, Mmp-9 is increased in the 30×30 cm cohort compared to the 50×50 cm cohort while no difference was noted for handling or between sexes. Data represent mean ± SEM with the number of animals indicated on legends and graphs by individual data points; *p ≤ 0.05, **p ≤ 0.01, ***, p ≤ 0.001, ****p ≤ 0.0001.

## Discussion

We report for the first time, to our knowledge, experimental condition- and sex-specific alterations of the BBB in the dorsal but mostly, ventral, HIP following NOR test exposure in adult mice. A large arena was associated with less time interacting with objects along with increased stress-related circulating CORT and opposing effects on BBB tight junction expressions in the ventral vs dorsal HIP (**Fig.7**). Conversely, handling improved recognition memory, reduced blood CORT and increased growth factor expression. Profound sex differences were noted not only after the NOR paradigm but even at baseline for the neurovascular genes analyzed in both HIP subregions. Finally, we identified sE-selectin and Mmp9 as blood vascular biomarkers affected by NOR testing that could help optimize experimental conditions when assessing recognition memory, to gain mechanistic insights in learning and memory processes or in the context of neurodegenerative or psychiatric diseases.

**Figure 7.**
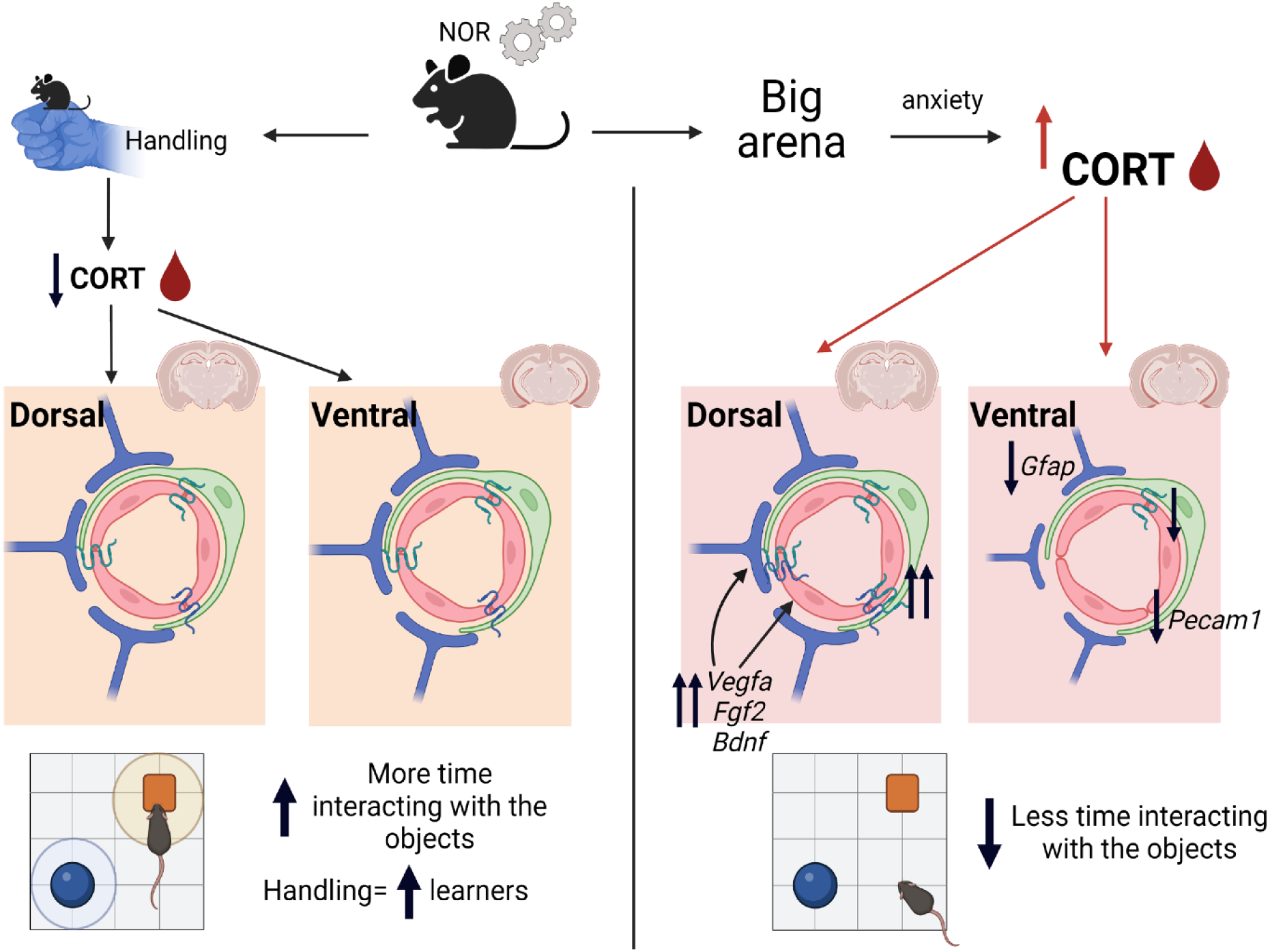
Summary of NOR-induced neurovascular changes in the dorsal vs ventral hippocampus. Mice that were subjected to the recognition memory test and were handled prior showed increased time spent with the objects and were overall considered better learners. At the biological level, they had lower circulating CORT and displayed no change in expression of BBB-related genes. Conversely, mice that experienced NOR in a bigger arena were characterized by anxious behaviors including less time spent interacting with the objects. Reflecting altered behaviors, blood CORT levels were higher, along with changes in several genes associated with the hippocampus BBB cells in a region-specific manner.

NOR memory testing is based on the natural tendency of rodents to explore novelty, hence, it is essential to reduce any stress or anxiety that may interfere with their desire to explore the arena and then, the objects. Handling has been recommended in previous publications not only for NOR (Leger et al., 2013; Lueptow, 2017) but also for behavioral tests associated with anxiety and stress responses (Hurst and West, 2010). Here, only NOR task was performed but it would be interesting to add an object location component to investigate further impact of short-term vs long-term memory formation on the BBB while avoiding inherent stress-induced by other paradigms such as fear conditioning or the Morris Water Maze (Vogel-Ciernia and Wood, 2014). Despite its increased use over the years, the underlying biological mechanisms supporting NOR learning processes in rodents have yet to be clearly defined including for the role of the HIP (Cohen and Stackman, 2015). Lack of consistency between NOR parameters across labs contributes to the debate (Antunes and Biala, 2012; Cohen and Stackman, 2015). However, as shown here, variation in experimental conditions greatly impact not only behavioral performances but also BBB biology, which could affect neuronal encoding considering the important role for the neurovascular unit in proper brain function. To improve rigor and reproducibility, it was recently proposed to favor 3D-printed objects (Inayat et al., 2021) which could be considered in future studies.

Exposure to a novel environment and learning context is stressful for rodents and associated with activation of the autonomic nervous system and hypothalamic-pituitary-adrenal (HPA) axis (Menard et al., 2017a). HPA activation lead to increased circulating CORT which can cross the BBB to activate steroid hormone receptors found in several brain regions and mediating nearly every aspect of brain function, including cognition, learning and memory (McEwen et al., 2015). In fact, injection of CORT immediately after a 3-min training trial can enhance NOR 24h retention performance in rats not previously habituated to the experimental context (Okuda et al., 2004). This effect is not observed if the animals received extensive prior habituation reducing emotional arousal during training (Okuda et al., 2004). CORT can bind to two types of receptors in the brain, mineralocorticoid or glucocorticoid, with the first being enriched in the lateral septum and HIP and the latter more widely distributed (Reul and de Kloet, 1985). Mineralocorticoid receptor *Nr3c2* is most highly expressed in astrocytes followed by endothelial cells (Zhang et al., 2014). As for glucocorticoid receptor *Nr3c1* highest expression is found in astrocytes, oligodendrocyte progenitor cells and endothelial cells (Zhang et al., 2014). Enriched transcriptomic profiles in cell types associated with the neurovascular unit – namely astrocytes and endothelial cells - may explain the changes we observed in BBB-related gene expression along with circulating CORT following NOR testing.

Sex hormones, including estrogens, progesterone, and androgens, affect emotions and cognition, therefore contributing to sex differences in behaviors (ter Horst et al., 2012; Dion-Albert et al., 2022a). Females respond differently to stress according to the phase of the estrous cycle with proestrus females being less anxious than their counterparts in other phases (ter Horst et al., 2012). Notably, binding capacity of HIP CORT receptors is higher in female rats, while the affinity is higher in males (Turner and Weaver, 1985). This dimorphism is driven by mineralocorticoid, and not glucocorticoid, receptors suggesting that under low CORT levels this stress response is less activated (Turner, 1997). Though, under chronic stress, the sex-specific patterns of mineralocorticoids and glucocorticoids change (ter Horst et al., 2012) and the estrus cycle can be disrupted (Herzog et al., 2009). In our study, we did not observe significant differences in estrous cycle phase between mice with high vs low NOR ratio (data not shown), however, we found profound sex differences in the expression of BBB-related genes in both dorsal and ventral HIP. Estrogen can cross the BBB and be produced endogenously in the brain. Nevertheless, its role on vascular-related functions remains poorly understood despite expression of both estrogen receptor subtypes, alpha and beta, on endothelial and vascular smooth muscle cells (Dion-Albert et al., 2022a). The brain endothelial and vascular smooth muscle cells also express androgen receptors that can modulate cerebrovascular reactivity, angiogenesis, and inflammatory processes (Abi-Ghanem et al., 2020). Both androgens and estrogens hormones, can cross the BBB by transmembrane diffusion in a bidirectional manner due to their small size and lipid solubility (Banks et al., 2020) increasing the complexity to define their role in behavior-driven changes in neurovascular function.

Growth factors can modulate BBB integrity and function. Astrocyte-derived Vegf-a drives BBB disruption in CNS inflammatory diseases (Argaw et al., 2012) and stress-induced depression (Matsuno et al., 2022). Bdnf is essential to promote persistence of long-term memory storage (Bekinschtein et al., 2008) and it can be transported across the BBB (Pan et al., 1998). In fact, blocking BDNF signaling after retrieval impairs object memory reconsolidation (Radiske et al., 2017). As for Fgf2, it is actively involved in the establishment and maintenance of the BBB (Reuss et al., 2003). It would be interesting to individually modulate these growth factors level in the dorsal and ventral HIP subregions in future studies to decipher their specific role in memory-induced changes in BBB properties. Nonetheless, it is crucial to mention that even if the dorsal HIP is generally linked with memory and spatial navigation whilst the ventral subregion has been associated with anxiety-related behaviors, it remains a topic of debate (Strange et al., 2014).

We recently identified sE-selectin as a vascular biomarker of stress responses and mood disorders (Dion-Albert et al., 2022b). This supports a clinical study associating higher circulating sE-selectin with microvascular dysfunction and low cognitive performances (Rensma et al., 2020). These reports are in line with the current finding that performing NOR in an anxious environment reduces time spent with the objects along with elevated blood sE-selectin. Further, we observed changes in Mmp9 which is an endopeptidase with various function in the brain including tissue formation, neurogenesis, and angiogenesis (Rempe et al., 2016). It is generally linked with BBB opening and hyperpermeability in disease conditions (Aoki et al., 2002; Shigemori et al., 2006; McMillin et al., 2015). Intriguingly, it is required for HIP late-phase long-term potentiation and memory formation (Nagy et al., 2006). Here, we show that circulating Mmp9 is reduced when mice are performing NOR in a smaller arena or when they are handled prior. Accordingly, elevated blood Mmp9 was associated with neuropsychiatric disorders often characterized by cognitive deficits (Beroun et al., 2019).

NOR memory becomes impaired in 18-25-month-old rodents (Benoit et al., 2011; Menard et al., 2013). In fact, novel object preference starts to decrease earlier (9-12 months) when temporal dynamics of exploration time are investigated (Traschutz et al., 2018). Here we focused strictly on NOR ratio and total interaction time with the objects. It would be interesting to explore further how the BBB reacts at various time points using *in vivo* imaging techniques (Lee et al., 2018). Indeed, under chronic stress, changes in BBB permeability are associated with altered levels of growth factor Vegfa and tight junction Cldn5 in line with our findings (Menard et al., 2017b; Lee et al., 2018; Dion-Albert et al., 2022b; Matsuno et al., 2022), suggesting that learning and memory processes may also modify BBB properties. Methylation at the *Cldn5* gene promoter represses its expression in mice (Dudek et al., 2020) and differential methylation patterns of human *CLDN5* were recently associated with cognitive decline (Huls et al., 2022). BBB breakdown is an early event in the aging human brain worsening mild cognitive impairment (Montagne et al., 2015). A large study exploring the genetics of memory function in ∼40,000 older individuals identified an association between *CLDN5* polymorphisms and verbal declarative memory performance (Debette et al., 2015). To summarize, it will be important to further explore the neurovascular contribution to learning and memory processes, in various emotional and environmental contexts, whilst considering sex and gender as biological variable throughout lifespan.

## Acknowledgements

This work was supported by the Natural Sciences and Engineering Research Council of Canada (Discovery Grant to C.M., undergraduate student research award for A.T.), Fonds de recherche du Quebec – Health (PhD scholarship to L.D.A., junior 1/junior 2 salary awards to C.M.), and the Canadian Institutes for Health Research (PhD scholarship to L.D.A.). The authors would like to thank Sam E.J. Paton for editing this manuscript.

---

**Supplementary Table 1.** Primers Table

| **Gene** | **Ref Seq #** | **Assay ID** | **Forward primer** | **Reverse primer** |
| --- | --- | --- | --- | --- |
| *Gapdh* | NM_008084(1) | Mm.PT.39a.1 | Exon location 2-3 |  |
| *Bdnf* | NM_001048139(1) | Mm.PT.58.8157970 | Exon location 2-5 |  |
| *Fgf2* | NM_008006(1) | Mm.PT.56a.5129235 | Exon location 1-3 |  |
| *Vegfa* | NM_001025250(3) | Mm.PT.58.14200306 | Exon location 1-2 |  |
| *Cldn5* | NM_013805(1) | Mm.PT.58.33394738.g | Exon location 1-1 |  |
| *Ocln* | NM_008756(1) | Mm.PT.58.42749240 | Exon location 7-9 |  |
| *Tjp1* | NM_001163574(2) | Mm.PT.58.12952721 | Exon location 25-26 |  |
| *Gfap* | NM_010277(1) | Mm.PT.58.31297710 | Exon location 6-9 |  |
| *Pecam1* | NM_001032378(2) | Mm.PT.58.43167370 | Exon location 7-8 |  |

